# HighFold-Lasso: Topology-Guided Prediction of Lasso Peptide Structures Using AlphaFold3

**DOI:** 10.64898/2026.10.14.759041

**Authors:** Sen Cao, Chengyun Zhang, Hanjie Yang, Yutong Ge, Hongli Hou, Ning Zhu, Yaling Wu, Jisong Mo, Juan Guo, Chongyang Li, Jingjing Guo, Hongliang Duan

**Author notes:** These authors contributed equally to this work (Sen Cao, Chengyun Zhang, Hanjie Yang). Author to whom correspondence should be addressed. Hongliang Duan.

## Abstract

Lasso peptides are a class of ribosomally synthesized and post-translationally modified peptides (RiPPs) characterized by a mechanically interlocked topology, in which the C-terminal tail is threaded through an N-terminal macrolactam ring formed by an isopeptide bond. This unique topology often confers high thermal stability and resistance to proteolytic degradation. However, the isopeptide linkage, threaded topology, and class-specific disulfide connectivity make lasso peptide structures difficult to reconstruct reliably using general-purpose structure prediction models. Here, we developed HighFold-Lasso, which combines custom covalent connectivity with lasso scaffold-guided adaptive low-noise initialization to enable conditional structure generation guided by lasso conformational priors. On a test set of 52 experimentally determined structures, HighFold-Lasso achieved a 100% lasso-topology success rate, with mean RMSD_Cα_ of 1.942 Å for the ring–loop region and 2.375 Å for the full peptide. The framework supports the modeling of the four classical lasso peptide classes (I–IV) and can represent noncanonical amino acids, providing computational support for structure–function studies and molecular engineering of lasso peptides.

## Introduction

Lasso peptides are a class of ribosomally synthesized and post-translationally modified peptides (RiPPs) characterized by a mechanically interlocked topology^1,2^. Their N-terminal α-amino group forms an isopeptide bond with the side-chain carboxyl group of an Asp or Glu residue, generating a macrolactam ring through which the C-terminal tail is threaded. Steric locks or disulfide bonds prevent the tail from unthreading, giving rise to the characteristic ring–loop– tail architecture^3,4^. Based on the number and connectivity of disulfide bonds, classical lasso peptides are divided into four classes (I–IV): class I contains two disulfide bonds, class II contains no disulfide bond, class III contains one ring–tail disulfide bond, and class IV contains one tail–tail disulfide bond^1,5,6^. More recently, lariocidin B, which contains a second intramolecular isopeptide bond, further expanded the known connectivity patterns of lasso peptides^7^. The conformational restriction imposed by the lasso topology often provides high thermal stability and resistance to proteolytic degradation, making lasso peptides attractive scaffolds for functional peptide development^1,8^. For example, lariocidin binds to a unique site on the bacterial ribosome and exhibits broad-spectrum antibacterial activity together with in vivo activity in a mouse infection model^7^. In addition, the incorporation of noncanonical amino acids and chemical modifications can further expand the chemical space of lasso peptides. The D-enantiomer of BI-32169, for example, shows substantially higher enzymatic stability than its natural L-form^9,10^. These developments further increase the need for structure modeling methods capable of handling both unusual topologies and nonstandard chemical components. Despite the thousands of candidate lasso peptide sequences identified through genome mining, only several dozen lasso peptides have experimentally determined three-dimensional structures^6,11^. One major obstacle is the difficulty in obtaining sufficient amounts of homogeneous mature peptides with the correct topology. In chemical synthesis, formation of the isopeptide bond does not necessarily produce a lasso conformation. Without appropriate preorganization, a peptide can form a branched-cyclic structure with the same covalent connectivity but without the required threading. Although the total synthesis of BI-32169 has been achieved using a multi-site preorganization strategy, this approach has not yet been successfully generalized to other lasso peptides^9^. Consistent with this difficulty, conformational sampling studies using molecular simulations have shown that free peptides have a low probability of adopting a cyclization-competent pre-lasso conformation and that this state can readily transition toward entropically favored unthreaded conformations^12,13^. Biosynthesis can promote formation of the lasso topology through precursor recognition, leader peptide cleavage, and cyclization, but it remains dependent on factors such as enzyme–substrate compatibility^14–16^. The resulting gap in experimentally determined structures limits our understanding of sequence–structure–function relationships and restricts structure-guided molecular engineering.

General-purpose structure prediction models, including AlphaFold, ESMFold, and Boltz, have greatly advanced computational structural biology^17–19^. However, accurately representing covalent connectivity while simultaneously recovering the threaded topology of lasso peptides remains challenging. Previous work showed that AlphaFold3 reproduced the lasso topology for approximately 79% (37/47) of PDB-deposited lasso peptides^6^. In contrast, among 40 RODEO-mined sequences with unknown experimental structures, only three predictions (7.5%) adopted a lasso-like topology. To address the limitations of general structure predictors, Juarez et al. developed LassoHTP, a computational framework based on scaffold construction and molecular mechanics modeling^6,20^. Ouyang et al. subsequently developed LassoPred with an annotator–constructor architecture. The annotator first predicts up to three possible ring–loop– tail annotations, after which the constructor generates the corresponding three-dimensional structures through scaffold construction, sequence mutation, and molecular mechanics optimization. However, the current version of LassoPred does not explicitly model disulfide-bond-defined scaffolds and does not support the general modeling of noncanonical residues. Its authors also identify disulfide-containing lasso peptides as a limitation of the current framework. Meanwhile, recent progress in cyclic peptide structure prediction has suggested that explicitly introducing specialized topological information can extend the applicability of state-of-the-art structure prediction models. Representative examples include HighFold1–3, AfCycDesign, and CyclicBoltz1^21–25^. In our recent work, HighFold4 further extended the handling of covalent connectivity in AlphaFold3, allowing user-defined inter-residue chemical connections to participate directly in structure prediction^26^. However, correct covalent connectivity does not guarantee a correct mechanically interlocked topology. A diffusion process initialized from high levels of random noise must still reconstruct the unusual threaded conformation. Moreover, the mechanically interlocked topology of a lasso peptide is not explicitly determined by its amino acid sequence alone. As a result, general structure prediction models may generate high-confidence structures that nevertheless fail to preserve the required threading relationship. Structure-conditioned diffusion provides a possible strategy for addressing this problem. Partial diffusion in RFdiffusion enables backbone conformations to be resampled around an existing fold, with the extent of structural variation controlled by the amount of added noise^27^. SiteAF3 uses a conditional diffusion process in which the receptor structure is fixed and binding-pocket or hotspot information can be introduced to guide complex structure generation^28^. More recently, true-coordinate injection experiments in Boltz-Perturb showed that when a trajectory is guided into the correct conformational region at an appropriate stage of denoising, the network can continue to recover the correct binding mode^29^. The outcome also depends on the remaining noise level, indicating that excessive perturbation can weaken the structural information provided by the injected coordinates. Together, these observations suggest that starting from a physically reasonable structure can reduce the conformational search space while allowing a generative model to resample local conformations without losing key structural features.

Based on this idea, we developed HighFold-Lasso, a lasso peptide structure prediction framework that explicitly defines the isopeptide bond and class-specific disulfide connectivity. A lasso scaffold with the target threading relationship is introduced as both a structural template and the starting structure for low-noise diffusion, enabling conditional generation around a valid lasso conformation. HighFold-Lasso supports the classical classes I–IV and can represent noncanonical residues through the Chemical Component Dictionary (CCD)^30^. We evaluate the model in terms of both structural accuracy and recovery of the lasso topology. In addition, predicted structures are processed using HighRelax^31^, our physics-based postprocessing framework for noncanonical amino acids and complex peptide topologies, to improve local geometry and stereochemical quality while preserving the overall conformation.

## Materials and Methods

### HighFold-Lasso Framework

HighFold-Lasso was developed based on the AlphaFold3 framework. By combining covalent connectivity encoding, lasso templates, and a lasso scaffold-guided adaptive low-noise initialization strategy, HighFold-Lasso introduces both chemical connectivity and threading priors into the structure generation process (Fig. 1A). The model inputs include the peptide sequence, a topologically valid lasso scaffold generated by LassoPred, and user-defined isopeptide and optional disulfide bond connections.

**Fig. 1.**
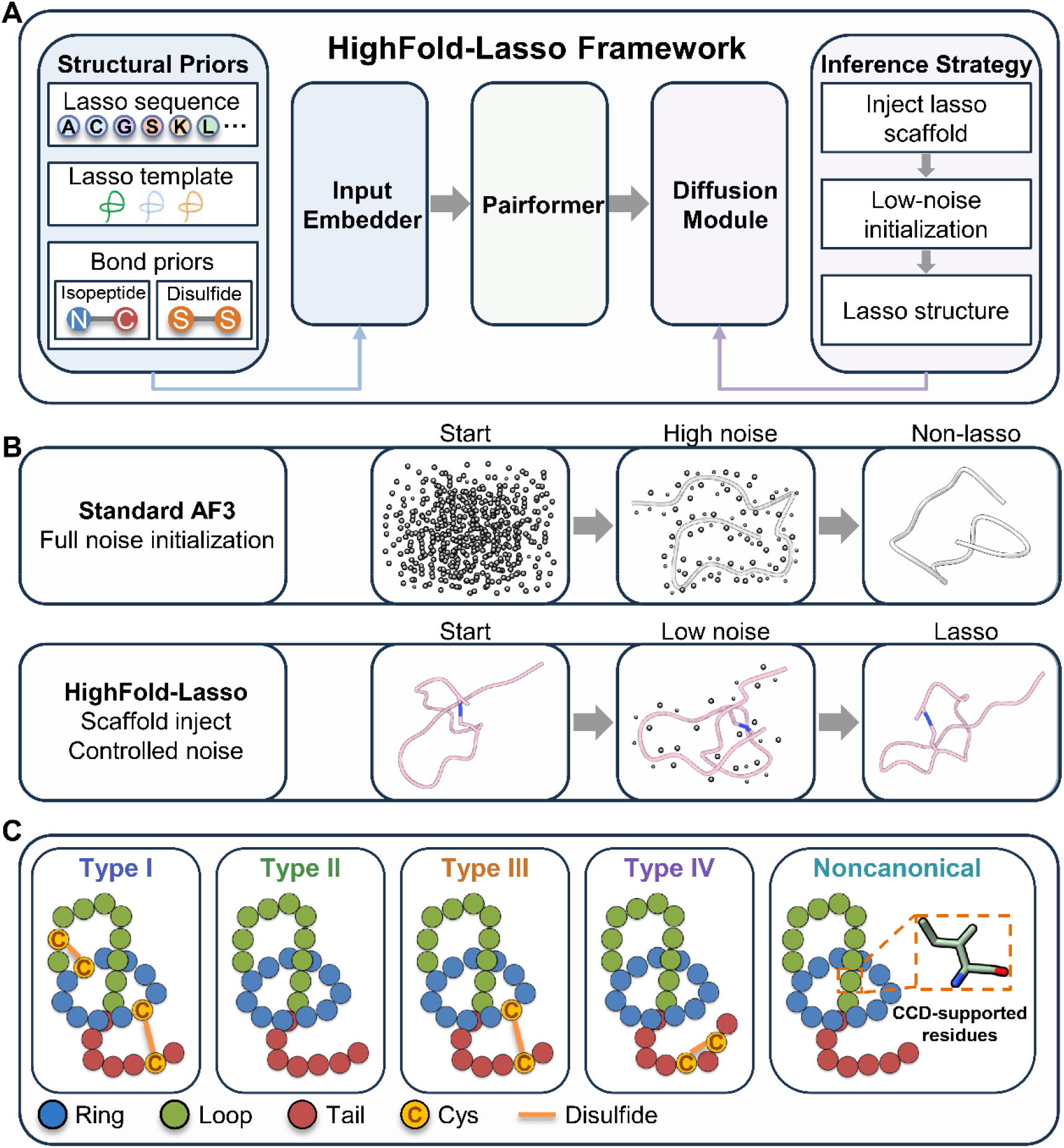
Overview of HighFold-Lasso. (A) Overall workflow of HighFold-Lasso, including model priors, lasso template input, and diffusion initialization. (B) Comparison between the standard AlphaFold3 diffusion process and the lasso scaffold-guided low-noise initialization strategy used in HighFold-Lasso. (C) Modeling support for classical class I–IV lasso peptides and systems containing noncanonical residues.

### Covalent Connectivity Representation

HighFold-Lasso explicitly incorporates inter-residue covalent connectivity during feature processing. The specified pairs of bonded atoms are first mapped to their corresponding token pairs, from which a covalent connectivity indicator matrix is constructed and encoded into the pair representation. This representation allows non-default covalent connections, such as isopeptide and disulfide bonds, to directly participate in subsequent structure generation rather than relying on the model to infer these connections implicitly from sequence or template information.

For lasso peptides, the isopeptide bond can be explicitly defined between the N-terminal nitrogen atom and the side-chain carbonyl carbon of the cyclizing Asp or Glu residue. Disulfide bonds can likewise be specified between the SG atoms of the corresponding Cys residues.

### Lasso Scaffold-Guided Adaptive Low-Noise Initialization

The lasso scaffold participates in structure generation through two complementary pathways. First, it is provided as a structural template to supply spatial information on the lasso conformation. Second, the same scaffold is used to initialize the coordinates of the diffusion process.

Unlike the default AlphaFold3 diffusion process, which starts from random coordinates at a high-noise state, HighFold-Lasso initializes diffusion from a lasso conformation with the correct threading relationship and applies controlled coordinate noise before denoising (Fig. 1B). Because different lasso scaffolds show different levels of tolerance to coordinate perturbation, multiple noise levels are progressively tested for each initial scaffold. The highest noise level that preserves the threaded topology is then selected as the starting point for diffusion sampling.

The noise schedule is defined as:

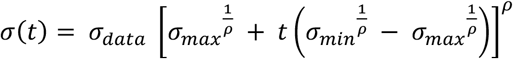

As *t* increases from 0 to 1, the noise level gradually decreases from the high-noise regime to the low-noise regime.

HighFold-Lasso controls the initial perturbation strength by adjusting the starting position *q* of the noise schedule. For a given *q*, the diffusion time points are defined as:

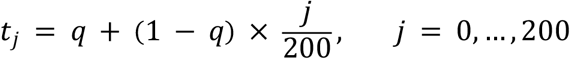

Thus, the strategy changes only the noise interval covered by the 200 reverse-diffusion updates, while keeping the number of diffusion updates and the core sampling procedure unchanged. For each starting position *q*, 201 noise-schedule points are generated and a fixed total of 200 reverse-diffusion updates are performed.

### Modeling of Multiple Lasso Peptide Classes and Noncanonical Residues

HighFold-Lasso supports structure modeling of the classical class I–IV lasso peptides as well as systems containing noncanonical amino acids (unAAs) (Fig. 1C). Different chemical topologies can be represented by explicitly defining the isopeptide bond together with optional disulfide bond connections. Noncanonical residues are represented using their corresponding components in the CCD, thereby extending the atomic representation beyond canonical amino acids.

### Isopeptide Bond Parameterization and Structure Optimization

Force field parameters for the isopeptide linkage were developed following the standard AMBER parameterization workflow (Fig. 2)^31–33^. First, according to the residue connectivity, one ACE and two NME capping groups were added to cap the exposed backbone termini and construct the dipeptide model (Step 1). Geometry optimization was then performed in the gas phase at the B3LYP/6-31G(d) level, followed by an HF/6-31G(d) calculation to obtain the electrostatic potential used for charge fitting (Step 2).

**Fig. 2.**
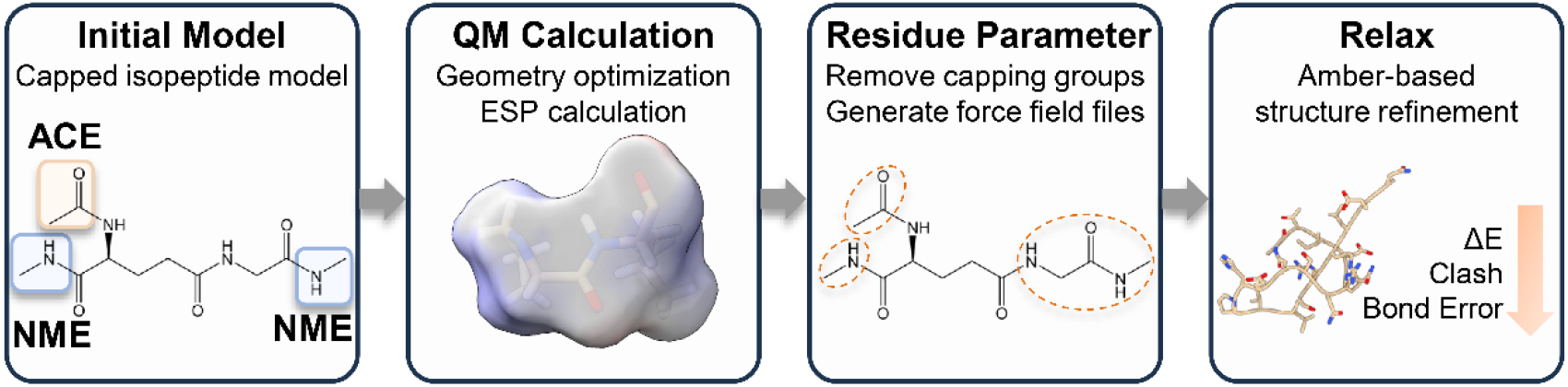
Workflow for isopeptide-linked residue parameterization and structure optimization. Using the GLU–GLY isopeptide model as an example, the workflow includes capped model construction, quantum chemical calculations, residue parameter generation, and subsequent structure optimization.

Electrostatic potential (ESP) data were extracted from the Gaussian output files using the espgen module in AmberTools, and atomic partial charges were derived by restrained electrostatic potential (RESP) fitting. After charge fitting, the capping groups and auxiliary residues were removed to extract the target residue parameters corresponding to the isopeptide-linked environment. For example, for the GLU–GLY model, GLY and the capping groups were removed, while the parameters of the ASP residue involved in the isopeptide linkage were retained. Missing force field terms were subsequently supplemented using the General Amber Force Field (GAFF). The final .prepi and .frcmod parameter files were generated, and atom names were standardized to match those in the corresponding PDB structures (Step 3).

The resulting parameters were integrated into HighRelax. Topology and coordinate files for the lasso peptides were generated using tleap in AmberTools, followed by energy minimization using OpenMM^34^. During minimization, harmonic positional restraints with a force constant of 10 kcal·mol^−1^·Å^−2^ were applied to all Cα atoms, and a maximum of 2000 minimization iterations was performed. This procedure corrects bond lengths at key covalent linkages, relieves steric clashes, and improves the physical plausibility of the predicted structures.

### Datasets

#### Experimentally Determined Lasso Peptide Dataset

The experimental structure benchmark was constructed starting from the 50 lasso peptide entries curated in the LassoPred benchmark. For complex structures, non-lasso binding partners were removed and only the lasso peptide chains were retained. Newly reported structures collected up to September 2026 were then added, resulting in a final dataset of 52 experimentally determined lasso peptide structures (Table S1). The dataset covers peptide lengths ranging from 14 to 33 amino acid residues and includes nine samples containing disulfide bonds and four samples containing noncanonical residues.

#### Genome-Mined Lasso Peptide Sequence Dataset

To evaluate the generalizability of HighFold-Lasso to genome-mined lasso peptide sequences, we constructed an extended test set from the 4,749 lasso peptide sequences curated by LassoPred from RODEO genome-mining results.11 A total of 100 sequences were selected by stratified random sampling with a fixed random seed, covering short (≤18 aa), medium-length (19–24 aa), and long (>24 aa) peptides (Table S2). To reduce potential overlap with experimentally characterized lasso peptides, sequences showing more than 80% global sequence identity to any sequence in the experimental structure dataset were excluded before sampling.

### Metrics

#### Root Mean Square Deviation Calculation

Root mean square deviation (RMSD) was used to evaluate structural differences between predicted and experimentally determined structures. For each sequence, the highest-confidence candidate among those satisfying the lasso conformation criteria was selected as the Top-1 prediction. If none of the candidates satisfied the criteria, the highest-confidence prediction was retained for structural deviation analysis.

Predicted and experimental structures were rigidly superimposed using the Kabsch algorithm. The evaluation metrics included Cα root mean square deviation(RMSD_Cα_), all-atom root mean square deviation(RMSD_all-atom_), and unAA root mean square deviation(RMSD_unAA_). For monomeric lasso peptides, RMSD_Cα_ and RMSD_all-atom_ were evaluated for the ring, ring–loop, and full-peptide regions after full-peptide superposition. For structures containing noncanonical amino acids, the noncanonical residue and neighboring atoms within a 10 Å radius were extracted for local structural superposition and RMSD calculation.

#### Structure Assessment

MolProbity was used to evaluate the stereochemical quality of both predicted and optimized structures. The assessment included steric clashes, geometric abnormalities, and overall structural quality, providing a complementary measure of the physical quality of predicted lasso peptide structures.

#### Lasso Conformation Assessment

The threading state was determined using a geometry-based criterion that evaluates whether the C-terminal peptide chain passes through the specified ring region. This criterion describes only the threading relationship and does not by itself define a complete lasso conformation. A predicted structure was classified as a valid lasso conformation only when two conditions were simultaneously satisfied: the C-terminal chain adopted the required threaded state, and the distance between the N-terminal nitrogen atom and the side-chain carbonyl carbon of the isopeptide-forming Asp or Glu residue was less than 4.0 Å.

## Results

### Benchmarking HighFold-Lasso on experimentally determined lasso peptide structures

We systematically compared the prediction performance of AlphaFold3, Boltz-2, LassoPred, ESMFold2, HighFold4, and HighFold-Lasso on a test set containing 52 experimentally determined lasso peptide monomer structures (Fig. 3A,B). AlphaFold3 achieved a mean RMSD_Cα_ of 2.72 Å and a lasso-conformation success rate of 67.3%. HighFold4 achieved a mean RMSD_Cα_ of 2.666 Å. The introduction of explicit covalent connectivity increased its lasso-conformation success rate to 75.0%, but it still failed to cover all samples. Boltz-2 and ESMFold2 showed mean RMSD_Cα_ of 3.60 and 5.40 Å, with success rates of 46.2% and 3.8%, respectively. LassoPred generated lasso-shaped structures for all samples, but its mean RMSD_Cα_ remained 3.50 Å. Overall, both current state-of-the-art general-purpose structure prediction models and lasso peptide-specific methods show some ability to model lasso peptides, but it remains difficult to simultaneously achieve robust recovery of the lasso conformation and high structural accuracy.

**Fig. 3.**
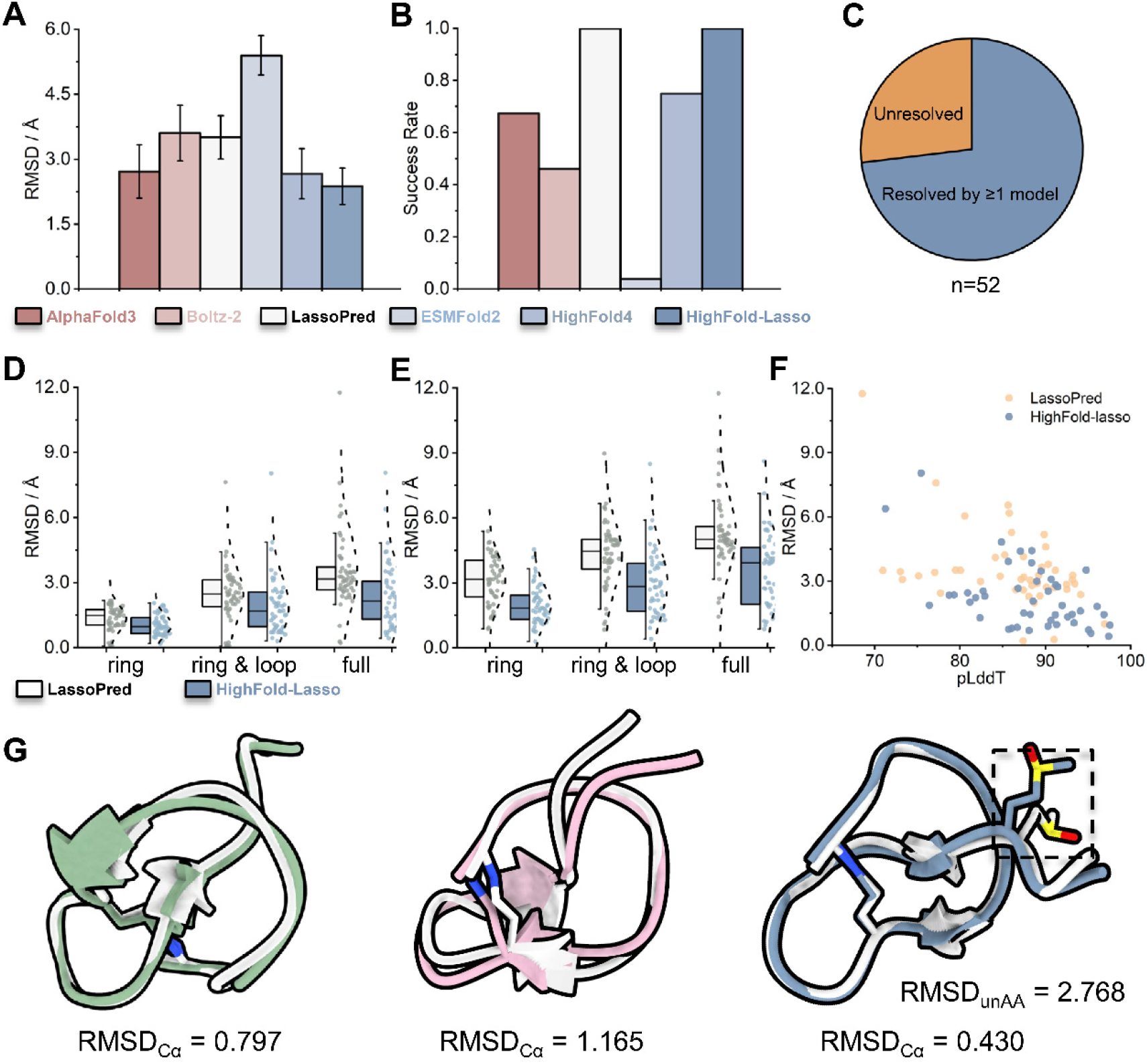
Prediction performance of different methods on the lasso peptide monomer dataset. (A) Full-peptide RMSD_Cα_; error bars indicate 95% confidence intervals. (B) Lasso-conformation success rate. (C) Combined sample coverage of AlphaFold3, Boltz-2, and ESMFold2. (D) Distributions of RMSD_Cα_ for the ring, ring–loop, and full-peptide regions after full-peptide superposition. (E) Distributions of RMSD_all-atom_ for the ring, ring–loop, and full-peptide regions after full-peptide superposition. (F) Correlation between pLDDT and RMSD_Cα_. (G) Structural superposition of predicted and experimental structures for 5GVO, 2N5C, and 5D9E; the local environment of the noncanonical residue is highlighted for 5D9E. Experimental structures are shown in gray, whereas predicted structures are shown in green, pink, and blue.

When the predictions from AlphaFold3, Boltz-2, and ESMFold2 were combined, the three general-purpose models together covered only 73.1% of the test set, leaving 26.9% of the samples without any structure satisfying the lasso-conformation criteria (Fig. 3C). Notably, AlphaFold3 predictions that failed to recover the lasso topology still showed relatively high model confidence, with a mean pLDDT of 81.53. This indicates that model confidence alone does not reliably reflect whether the mechanically interlocked topology of a lasso peptide has been correctly recovered (Table S3). These results suggest that current general-purpose structure prediction models can recognize some known lasso conformations, but their ability to generalize across diverse lasso peptide structures remains limited.

To address these limitations, we developed HighFold-Lasso by integrating lasso scaffold priors, explicit covalent connectivity, and low-noise structural initialization. HighFold-Lasso achieved a 100% lasso-conformation success rate and the lowest mean RMSD_Cα_ of 2.375 Å among the evaluated methods (Fig. 3A,B). Because the tail region of lasso peptides is relatively flexible, we further evaluated structural deviations separately for the ring, ring–loop, and full-peptide regions. For RMSD_Cα_, HighFold-Lasso achieved mean values of 1.05, 1.94, and 2.38 Å for the three regions, respectively, representing reductions of 21.7%, 20.5%, and 32.2% compared with the corresponding LassoPred values of 1.35, 2.44, and 3.51 Å. The corresponding median RMSD_Cα_ values decreased from 1.48, 2.48, and 3.16 Å for LassoPred to 0.96, 1.69, and 2.15 Å for HighFold-Lasso. In paired comparisons, HighFold-Lasso produced lower RMSD_Cα_ values for 69.2%, 65.4%, and 76.9% of the samples in the ring, ring–loop, and full-peptide regions, respectively (Fig. 3D).

At the all-atom level, HighFold-Lasso achieved mean RMSD_all-atom_ values of 1.98, 2.98, and 3.55 Å for the ring, ring–loop, and full-peptide regions, respectively. These values were 36.6%, 30.6%, and 32.8% lower than those obtained with LassoPred (3.12, 4.30, and 5.29 Å, respectively). For each of the three structural regions, 90.4% of the paired samples showed lower RMSD_all-atom_ values with HighFold-Lasso (Fig. 3E). Using an RMSD threshold of 2 Å, HighFold-Lasso achieved RMSD_Cα_ < 2 Å for 51, 31, and 24 samples in the ring, ring–loop, and full-peptide regions, respectively, compared with 44, 14, and 4 samples for LassoPred. At the all-atom level, the corresponding numbers increased from 10, 5, and 1 for LassoPred to 31, 16, and 13 for HighFold-Lasso. Two-sided paired Wilcoxon signed-rank tests showed significant differences in all six RMSD comparisons (P < 0.001). These results indicate that the improvement achieved by HighFold-Lasso is not driven by a small subset of samples. Instead, lasso scaffold-guided diffusion consistently improves structural agreement across different regions, at both the backbone and all-atom levels. Detailed results are provided in Tables S4 and S5.

To evaluate the consistency between model confidence and actual structural accuracy, we used AF3Score^35^ to score structures generated by LassoPred and HighFold-Lasso under a unified framework (Fig. 3F). AF3Score is an AlphaFold3-based structure scoring method that bypasses diffusion-based structure generation and directly feeds the input coordinates into the confidence module. HighFold-Lasso showed a negative correlation between pLDDT and RMSD_Cα_ (Pearson r = −0.507), stronger than that observed for LassoPred (Pearson r = −0.433). Under the high-confidence condition of pLDDT > 90, 15 HighFold-Lasso predictions simultaneously achieved RMSD_Cα_ < 2 Å, compared with only two LassoPred predictions. These results indicate a closer agreement between model confidence and structural accuracy for HighFold-Lasso. Detailed results are provided in Table S6.

We further evaluated the ability of different methods to generate lasso conformations for 100 genome-mined candidate lasso peptide sequences without experimentally determined three-dimensional structures. Because reference structures were unavailable for these sequences, this analysis focused only on whether the generated structures satisfied the predefined lasso-conformation criteria. AlphaFold3 generated valid lasso conformations for only 4% of the sequences, whereas both LassoPred and HighFold-Lasso achieved 100% (Table 1). These results suggest that scaffold-guided approaches can robustly generate structures that preserve the required lasso conformation across diverse genome-mined sequences.

**Table 1.** Lasso-conformation success rates of different methods on genome-mined lasso peptide sequences.

| Model | Success rate |
| --- | --- |
| AlphaFold3 | 4% |
| LassoPred | 100% |
| HighFold-Lasso | 100% |

Finally, representative examples were used to illustrate the structural accuracy of HighFold-Lasso. The predicted structures of sphaericin (PDB ID: 5GVO), chaxapeptin (PDB ID: 2N5C), and caulosegnin II (PDB ID: 5D9E) achieved RMSDCα of 0.797, 1.165, and 0.430 Å, respectively. Notably, caulosegnin II contains the modified residue SME. These examples suggest that HighFold-Lasso can closely reproduce experimentally determined lasso peptide conformations while preserving the mechanically interlocked topology and reducing structural deviations, and can be extended to lasso peptide systems containing nonstandard modified residues.

### Performance across classical lasso peptide classes

To evaluate the applicability of HighFold-Lasso to different lasso peptide topologies, we grouped the test set according to the classical classes I–IV. These classes differ in the number and connectivity patterns of disulfide bonds. Compared with LassoPred, HighFold-Lasso achieved lower structural deviations across all four classes. The mean RMSD_Cα_ values for classes I, II, III, and IV decreased from 4.24, 3.24, 4.23, and 5.75 Å with LassoPred to 1.88, 2.43, 1.05, and 2.56 Å with HighFold-Lasso, corresponding to reductions of 55.7%, 24.9%, 75.3%, and 55.4%, respectively (Fig. 4A). Similarly, the mean RMSD_all-atom_ values decreased from 5.94, 5.09, 5.44, and 6.76 Å to 2.99, 3.63, 1.99, and 3.38 Å, representing reductions of 49.7%, 28.6%, 63.5%, and 50.1%, respectively (Fig. 4B). Notably, improvements were also observed for class II lasso peptides, which contain no disulfide bonds, indicating that the performance gain cannot be attributed solely to the introduction of disulfide connectivity.

**Fig. 4.**
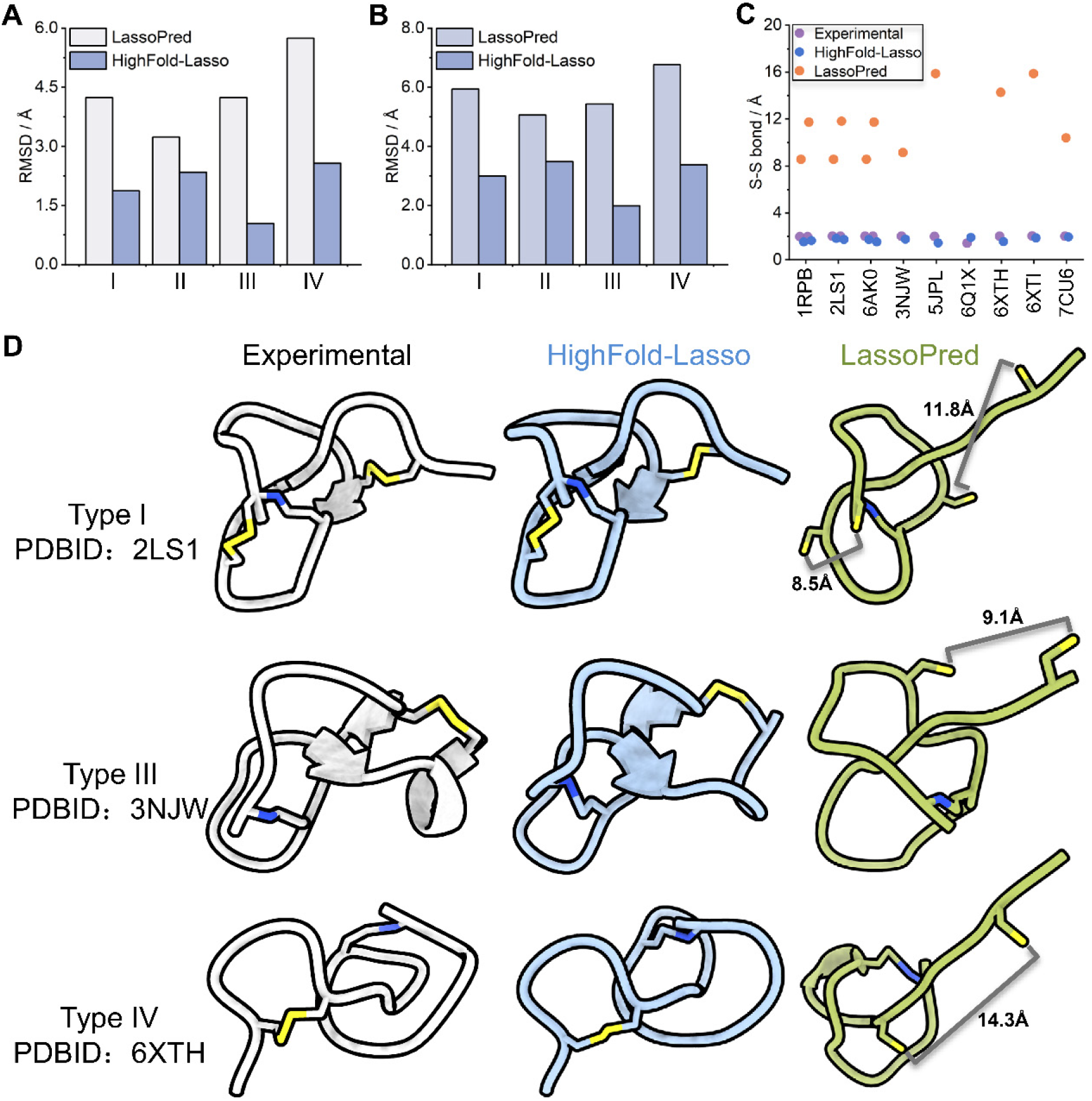
Prediction performance across different classes of lasso peptides. (A,B) Comparison of RMSD_Cα_ and RMSD_all-atom_ across class I–IV lasso peptides. 7CU6 contains a noncanonical disulfide topology and was therefore excluded from the standard class I–IV stratified analysis. (C) Distribution of disulfide bond lengths in predicted and experimental structures. (D) Representative structural comparisons for the class I peptide 2LS1, class III peptide 3NJW, and class IV peptide 6XTH.

To evaluate the effect of explicit covalent connectivity on the recovery of more complex lasso topologies, we further analyzed the S–S distances between the target Cys residues in the experimental and predicted structures (Fig. 4C). In the initial structures generated by LassoPred, the mean distance between the target cysteine sulfur atoms was 12.21 ± 3.63 Å. HighFold-Lasso reduced this distance to 1.70 ± 0.17 Å. Although this value was shorter than the experimental reference of 1.99 ± 0.12 Å and therefore required further bond-length correction, the result indicates that explicit connectivity information can drive spatially separated residues toward the intended covalent connection. Importantly, the initial lasso scaffolds provided to HighFold-Lasso did not explicitly contain disulfide bond connectivity. Thus, the observed rearrangement reflects the ability of the model to respond to newly introduced chemical connectivity while preserving the predefined threaded conformation.

To distinguish the contribution of the initial scaffold prior from subsequent conformational remodeling, we performed paired comparisons between the input scaffolds and the corresponding HighFold-Lasso outputs (Table S7). Independent geometric assessment confirmed that all input and output structures retained the lasso conformation. The mean RMSD_Cα_ between the input and output structures were 2.76 ± 1.58, 2.99 ± 1.50, and 3.23 ± 1.50 Å for the ring, ring–loop, and full-peptide regions, respectively. These deviations indicate that the output structures were not simple copies of the initial scaffolds. Together with the improved agreement with the experimental structures described above, these results support that HighFold-Lasso preserves the lasso topology of the input scaffold while further remodeling its conformation toward the experimentally observed state.

Representative structures further illustrate these differences (Fig. 4D). For the class I peptide 2LS1, class III peptide 3NJW, and class IV peptide 6XTH, HighFold-Lasso recovered both the lasso conformation and the corresponding disulfide connectivity. In contrast, LassoPred does not provide explicit control over disulfide bond connectivity in its current implementation. These results suggest that lasso scaffold-guided conditional generation can accommodate different disulfide connectivity patterns, providing a unified framework for modeling classical class I–IV lasso peptides and a suitable structural basis for subsequent local geometry optimization.

### Ablation Study

To investigate the contribution of individual components in HighFold-Lasso, we separately removed the explicit isopeptide connectivity, template features, and low-noise structural initialization. Their effects were evaluated in terms of overall structural accuracy, lasso-conformation success rate, and local isopeptide geometry.

We first compared the overall structural accuracy under different ablation settings. The full model achieved a mean RMSD_Cα_ of 2.38 Å. Removing the explicit isopeptide connectivity, template features, or low-noise structural initialization increased the mean RMSD_Cα_ to 2.56, 2.70, and 2.81 Å, corresponding to increases of 7.9%, 13.9%, and 18.4%, respectively (Fig. 5A). These results indicate that all three components contribute to structural accuracy, with removal of low-noise structural initialization producing the largest loss in overall conformational accuracy.

**Fig. 5.**
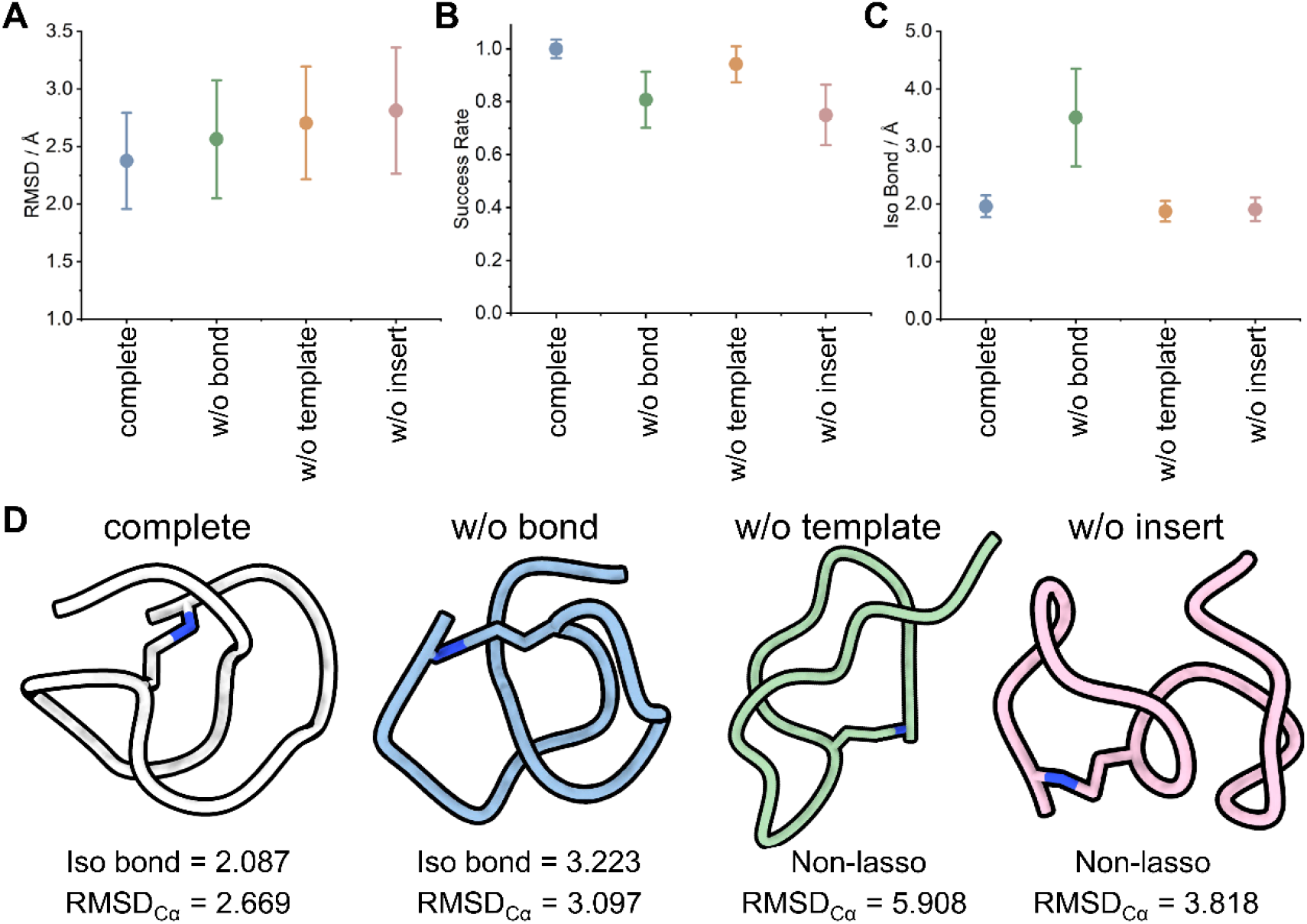
Ablation analysis of HighFold-Lasso. (A) Full-peptide RMSD_Cα_; error bars indicate 95% confidence intervals. (B) Lasso-conformation success rate; error bars indicate 95% confidence intervals. (C) Distance between the isopeptide-forming atoms; error bars indicate 95% confidence intervals. (D) Representative structures of 7ZWJ under different ablation settings.

We next examined the contribution of each component to recovery of the mechanically interlocked topology. The full HighFold-Lasso model generated valid lasso conformations for all test samples. The success rates decreased to 80.8%, 94.2%, and 75.0% after removing the isopeptide connectivity, template features, and low-noise structural initialization, respectively (Fig. 5B). The largest decrease was observed when low-noise structural initialization was removed, indicating that structure initialization plays a particularly important role in preserving the lasso conformation during diffusion sampling.

We further evaluated the effects of these components on local isopeptide geometry. The mean distance between the isopeptide-forming atoms was 1.96 Å for the full model. This distance increased to 3.50 Å when the explicit isopeptide connectivity was removed, whereas removal of the template features or low-noise initialization resulted in mean distances of 1.87 and 1.91 Å, respectively (Fig. 5C). These results indicate that explicit isopeptide connectivity primarily contributes to maintaining the local spatial relationship required for formation of the isopeptide linkage.

The representative example 7ZWJ further illustrates the distinct roles of these components (Fig. 5D). The full model retained the lasso conformation, with an RMSD_Cα_ of 2.669 Å and an isopeptide-forming atom distance of 2.087 Å. After removal of the explicit isopeptide connectivity, the peptide still adopted a lasso conformation, but the corresponding atom distance increased to 3.223 Å and the RMSD_Cα_ increased to 3.097 Å. In contrast, removal of either the template features or low-noise structural initialization resulted in loss of the lasso topology, with RMSD_Cα_ of 5.908 and 3.818 Å, respectively.

Together, these results suggest complementary roles for explicit covalent connectivity, template features, and low-noise structural initialization. Explicit covalent connectivity primarily maintains the local geometry associated with the intended chemical linkage, whereas template information and low-noise initialization contribute to recovery and preservation of the global lasso conformation.

### Physics-Based Optimization of HighFold-Lasso Predictions

To improve the physical plausibility of HighFold-Lasso predictions, we developed AMBER-compatible force field parameters for the characteristic isopeptide linkage of lasso peptides and incorporated them into HighRelax for energy minimization.

We first evaluated structural quality before and after relaxation using MolProbity. As shown in Fig. 6A, the mean MolProbity score decreased from 3.17 before relaxation to 1.86 after HighRelax, indicating a substantial improvement in overall stereochemical quality. Analysis of atomic steric clashes showed an even more pronounced improvement (Fig. 6B). The mean clash score decreased from 66.23 to 0.52 after relaxation, demonstrating that energy minimization effectively removes atomic overlaps and unfavorable short-range contacts and thereby improves the local physical quality of the predicted structures.

**Fig. 6.**
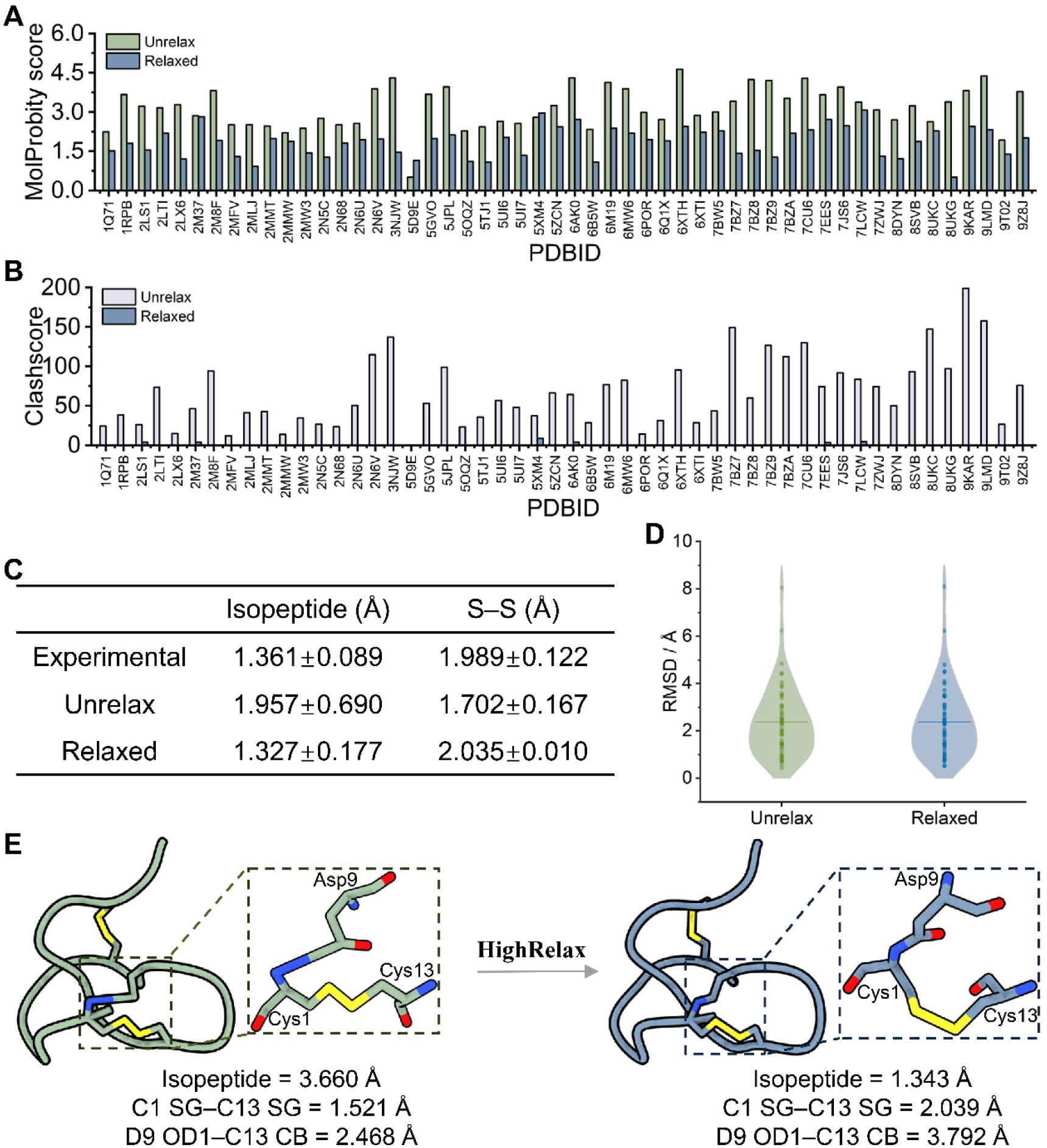
Effects of HighRelax on HighFold-Lasso predictions. (A) MolProbity scores before and after relaxation. (B) Clash scores before and after relaxation. (C) Isopeptide and disulfide bond lengths before and after relaxation compared with experimental references. (D) RMSD_Cα_ distributions before and after relaxation. (E) Representative structure of 1RPB showing local geometric changes and interatomic distances before and after HighRelax.

In addition to steric clashes, we examined the geometries of the characteristic covalent connections in lasso peptides (Fig. 6C). In the experimental structures, the mean isopeptide and disulfide bond lengths were 1.36 ± 0.09 Å and 1.99 ± 0.12 Å, respectively. Before relaxation, the corresponding distances in the predicted structures were 1.96 ± 0.69 Å and 1.70 ± 0.17 Å. After HighRelax, these values were adjusted to 1.33 ± 0.18 Å and 2.04 ± 0.01 Å, respectively, bringing both distributions closer to their experimental reference values. These results indicate that force field optimization incorporating explicit isopeptide parameters can correct residual local geometric deviations in covalent connections after diffusion-based structure generation. To determine whether HighRelax affected the overall structural accuracy already achieved by HighFold-Lasso, we further compared RMSD_Cα_ values relative to the experimental structures before and after relaxation (Fig. 6D). The mean RMSD_Cα_ was 2.375 Å before relaxation and 2.374 Å after relaxation, remaining essentially unchanged. Thus, HighRelax primarily improves local atomic arrangements, covalent bond geometry, and steric contacts without substantially altering the overall mechanically interlocked conformation predicted by HighFold-Lasso. This behavior is consistent with the intended role of HighRelax as a physics-based postprocessing procedure that improves local structural quality while preserving the global fold.

A representative example is shown for 1RPB (Fig. 6E). Before relaxation, the distance between the isopeptide-forming atoms was 3.660 Å, and the Cys1–Cys13 S–S distance was 1.521 Å. An unfavorable short contact of 2.468 Å was also observed between Asp9 OD1 and Cys13 CB. After HighRelax, the isopeptide bond distance was adjusted to 1.343 Å, the S–S distance was restored to 2.039 Å, and the D9 OD1–C13 CB distance increased to 3.792 Å. These changes directly illustrate the ability of HighRelax to correct covalent bond geometries and unfavorable local atomic arrangements, thereby improving the local geometric quality and physical plausibility of HighFold-Lasso predictions.

## Discussion

The major challenge in lasso peptide structure prediction is the correct formation and preservation of the characteristic threaded topology. General-purpose structure prediction models can recover the lasso conformation for some experimentally characterized peptides, but their performance decreases substantially for sequences lacking closely related structural information. Recent studies of lasso peptide folding have reached a similar conclusion. Molecular simulations show that spontaneous formation of threaded or pre-lasso conformations is rare, because formation of the constrained threaded state carries a substantial entropic penalty and competes with more accessible unthreaded conformations. This behavior is consistent with our observations: some predictions that failed to recover the correct lasso topology still received high confidence scores, indicating that high model confidence does not necessarily imply topological correctness. A related issue was noted in our previous HighFold3 study, where peptide topology was not uniquely determined by sequence alone, and the same sequence could adopt different structural forms depending on cyclization or processing conditions. Therefore, for lasso peptides, sequence information and general structural priors alone may be insufficient to reliably recover the mature mechanically interlocked conformation.

Based on this consideration, HighFold-Lasso introduces a lasso scaffold, explicit covalent connectivity, and structure-initialized low-noise diffusion so that structure generation starts from a conformation with the desired threading relationship rather than requiring the model to reconstruct the lasso topology from a random high-noise state. This strategy improved both the lasso-conformation success rate and structural accuracy across different peptide regions. Importantly, the initial lasso scaffolds did not explicitly contain disulfide connectivity. When explicit Cys–Cys connections were subsequently introduced, HighFold-Lasso produced corresponding spatial rearrangements of the cysteine residues. This observation indicates that the diffusion process can remodel relative residue positions in response to newly introduced chemical connectivity while retaining the pre-existing threaded topology. HighFold-Lasso can therefore be viewed as a form of lasso scaffold-guided conditional generation, in which the output preserves key topological information from the initial scaffold while incorporating sequence and chemical-connectivity information to generate a new conformation.

The ablation experiments further indicate that these structural priors act at different levels. Explicit covalent connectivity primarily promotes the local spatial arrangement required for the intended chemical linkage, whereas template features and structure initialization improve preservation of the global threading relationship. At the same time, the structures generated directly by diffusion still contained local geometric imperfections, including elongated isopeptide distances, shortened disulfide distances, and steric clashes. These observations highlight a distinction between recovering the correct topology and obtaining physically realistic local geometry. By incorporating dedicated isopeptide parameters, HighRelax effectively corrected these remaining local defects while leaving the overall lasso conformation nearly unchanged. This behavior is consistent with the original role of HighRelax as a physics-based postprocessing framework that improves local stereochemistry and atomic geometry without substantially perturbing the global fold.

The same general strategy may be transferable to other diffusion-based structure prediction frameworks that support structural conditioning or inference-time guidance. For example, Boltz-2 provides inference-time steering potentials that can guide diffusion toward user-specified structural features, while Protenix^36^ includes a training-free guidance module that applies geometric and physical potentials during diffusion sampling. Combining such guidance mechanisms with lasso scaffold initialization may help improve local physical plausibility during generation and reduce the amount of postprocessing required. However, whether these potentials can preserve a mechanically interlocked threading state while maintaining or improving structural accuracy remains to be systematically evaluated.

We also explored extending HighFold-Lasso toward protein–lasso peptide complex prediction. Recent coordinate-injection experiments in Boltz-Perturb provide a useful analogy. In protein– small-molecule systems, replacing intermediate diffusion coordinates with experimentally correct coordinates allowed subsequent denoising to recover the correct binding mode in some cases, whereas injection at earlier, higher-noise stages often returned to the default incorrect solution. These results suggest that at least part of the prediction error can arise from insufficient sampling and that an appropriate structural starting region can help guide the denoising trajectory toward a native-like conformational basin. Guided by this idea, we used coarse binding regions predicted by AlphaFold3 and introduced lasso peptides with valid threaded topologies during diffusion, while sampling their positions and orientations. For lasso peptides, however, interface sampling is more difficult because their relatively large size and topologically constrained conformation require coordinated changes in translation, rotation, and local peptide geometry. Increasing the diffusion noise can broaden the search over possible binding modes, but excessive noise can simultaneously disrupt the preformed threaded topology.

Our preliminary exploration suggests that near-native candidate poses can be sampled for many targets, but current confidence scores, conformational clustering, and energy-minimization-based filtering do not consistently rank these candidates among the top predictions. This limitation is conceptually consistent with Boltz-Perturb, where confidence scores were also reported to be poorly suited for distinguishing alternative sampled poses in some coordinate-injection experiments. Thus, a major remaining challenge is the development of reliable pose-ranking methods that do not depend on knowledge of the experimental structure, while simultaneously improving sampling efficiency. At present, HighFold-Lasso has therefore been systematically validated primarily for lasso peptide monomers. Extension to protein–lasso peptide complexes will likely require generation and scoring strategies that explicitly preserve the lasso topology during interface sampling and can reliably identify native-like binding modes.

Overall, HighFold-Lasso provides an integrated workflow for structure prediction and optimization of classical class I–IV lasso peptides. By combining lasso scaffold priors, user-defined covalent connectivity, and low-noise diffusion with CCD-based representation of noncanonical residues and physics-based local optimization, the framework addresses both the mechanically interlocked topology and the chemical complexity of lasso peptides. It therefore provides a computational basis for studying lasso peptide structure–function relationships and for the structural modeling of lasso peptides containing nonstandard residues.

## SUPPLEMENTARY DATA

Supplementary data are available online.

## Data availability

The experimentally determined lasso peptide structures used for benchmarking in this study are available in the Protein Data Bank (PDB), with their accession codes provided in Supplementary Table S1. The genome-mined sequences are provided in Supplementary Table S2. All other data generated or analysed during this study are included in this published article and its supplementary information files. Source data and related structure files are provided in the associated GitHub repository (https://github.com/babysensen/HighFold-Lasso).

## AUTHOR CONTRIBUTIONS

Sen Cao: Conceptualization, Data curation, Methodology, Formal analysis, Supervision, Writing – original draft. Chengyun Zhang: Methodology, Supervision. Hanjie Yang: Data curation, Methodology. Yutong Ge: Methodology. Hongli Hou: Methodology. Ning Zhu: Methodology. Yaling Wu: Methodology. Jisong Mo: Methodology. Juan Guo: Methodology. Chongyang Li: Methodology. Jingjing Guo: Methodology. Hongliang Duan: Conceptualization, Supervision, Funding acquisition. All authors contributed to the writing, review, and editing of the manuscript and approved the final version.

## Code availability

The source code and scripts used to implement HighFold-Lasso and reproduce the analyses in this study are publicly available in the HighFold-Lasso GitHub repository (https://github.com/babysensen/HighFold-Lasso).

## REFERENCES

1. Maksimov, M. O., Pan, S. J. & James Link, A. Lasso peptides: structure, function, biosynthesis, and engineering. Nat. Prod. Rep. 29, 996 (2012).

2. Montalbán-López, M. et al. New developments in RiPP discovery, enzymology and engineering. Nat. Prod. Rep. 38, 130–239 (2021).

3. Do, T. & Link, A. J. Protein Engineering in Ribosomally Synthesized and Post-translationally Modified Peptides (RiPPs). Biochemistry 62, 201–209 (2023).

4. Hegemann, J. D. Factors Governing the Thermal Stability of Lasso Peptides. ChemBioChem 21, 7–18 (2020).

5. Hegemann, J. D., Zimmermann, M., Xie, X. & Marahiel, M. A. Lasso Peptides: An Intriguing Class of Bacterial Natural Products. Acc. Chem. Res. 48, 1909–1919 (2015).

6. Ouyang, X. et al. LassoPred: a tool to predict the 3D structure of lasso peptides. Nat. Commun. 16, 5497 (2025).

7. Jangra, M. et al. A broad-spectrum lasso peptide antibiotic targeting the bacterial ribosome. Nature 640, 1022–1030 (2025).

8. Barrett, S. E. & Mitchell, D. A. Advances in lasso peptide discovery, biosynthesis, and function. Trends Genet. 40, 950–968 (2024).

9. Chen, M., Wang, S. & Yu, X. Cryptand-imidazolium supported total synthesis of the lasso peptide BI-32169 and its D-enantiomer. Chem. Commun. 55, 3323–3326 (2019).

10. Potterat, O. et al. BI-32169, a Bicyclic 19-Peptide with Strong Glucagon Receptor Antagonist Activity from Streptomyces sp. J. Nat. Prod. 67, 1528–1531 (2004).

11. Tietz, J. I. et al. A new genome-mining tool redefines the lasso peptide biosynthetic landscape. Nat. Chem. Biol. 13, 470–478 (2017).

12. Da Hora, G. C. A., Oh, M., Nguyen, J. D. M. & Swanson, J. M. J. One Descriptor to Fold Them All: Harnessing Intuition and Machine Learning to Identify Transferable Lasso Peptide Reaction Coordinates. J. Phys. Chem. B 128, 4063–4075 (2024).

13. Yin, S., Mi, X., Barrett, S. E., Mitchell, D. A. & Shukla, D. De novo Folding Mechanisms of Lasso Peptides. Preprint at 10.64898/2026.03.30.715466 (2026).

14. Arnison, P. G. et al. Ribosomally synthesized and post-translationally modified peptide natural products: overview and recommendations for a universal nomenclature. Nat Prod Rep 30, 108–160 (2013).

15. Kretsch, A. M. et al. Peptidase Activation by a Leader Peptide-Bound RiPP Recognition Element. Biochemistry 62, 956–967 (2023).

16. Yan, K. et al. Dissecting the Maturation Steps of the Lasso Peptide Microcin J25 in vitro. ChemBioChem 13, 1046–1052 (2012).

17. Abramson, J. et al. Accurate structure prediction of biomolecular interactions with AlphaFold 3. Nature 630, 493–500 (2024).

18. Wohlwend, J. et al. Boltz-1 Democratizing Biomolecular Interaction Modeling. 2024.11.19.624167 Preprint at 10.1101/2024.11.19.624167 (2024).

19. Lin, Z. et al. Evolutionary-scale prediction of atomic-level protein structure with a language model. Science 379, 1123–1130 (2023).

20. Juarez, R. J. et al. LassoHTP: A High-Throughput Computational Tool for Lasso Peptide Structure Construction and Modeling. J. Chem. Inf. Model. 63, 522–530 (2023).

21. Cao, S. et al. Accurate structure prediction of cyclic peptides containing unnatural amino acids using HighFold3. Brief. Bioinform. 26, bbaf488 (2025).

22. Zhang, C. et al. HighFold: accurately predicting structures of cyclic peptides and complexes with head-to-tail and disulfide bridge constraints. Brief. Bioinform. 25, bbae215 (2024).

23. Zhu, C. et al. Predicting the structures of cyclic peptides containing unnatural amino acids by HighFold2. Brief. Bioinform. 26, bbaf202 (2025).

24. Xie, X., Li, C. Z., Lee, J. S. & Kim, P. M. CyclicBoltz1, fast and accurately predicting structures of cyclic peptides and complexes containing non-canonical amino acids using AlphaFold 3 Framework. Preprint at 10.1101/2025.02.11.637752 (2025).

25. Rettie, S. A. et al. Cyclic peptide structure prediction and design using AlphaFold2. Nat. Commun. 16, 4730 (2025).

26. Zhang, C. et al. HighFold4: extending AlphaFold3 to accurate cyclic peptide conformation prediction via custom chemical connectivity. Brief. Bioinform. 27, bbag505 (2026).

27. Watson, J. L. et al. De novo design of protein structure and function with RFdiffusion. Nature 620, 1089–1100 (2023).

28. Tang, H. & Wang, J. Accurate site-specific folding via conditional diffusion based on AlphaFold3. Proc. Natl. Acad. Sci. 122, e2521048122 (2025).

29. Jung, H., Lee, B. & Cheng, A. C. Boltz-Perturb: Improving Diversity and Accuracy in Protein-Ligand Co-Folding through Training-Free Conditioning Perturbation. Preprint at 10.64898/2026.08.05.742877 (2026).

30. Westbrook, J. D. et al. The chemical component dictionary: complete descriptions of constituent molecules in experimentally determined 3D macromolecules in the Protein Data Bank. Bioinformatics 31, 1274–1278 (2015).

31. Cao, S. et al. HighRelax: Physics-Based Refinement of Deep Learning Protein Predictions with Noncanonical Amino Acids. J. Chem. Theory Comput. 22, 3093–3102 (2026).

32. Wang, J., Wolf, R. M., Caldwell, J. W., Kollman, P. A. & Case, D. A. Development and testing of a general amber force field. J. Comput. Chem. 25, 1157–1174 (2004).

33. Cornell, W. D., Cieplak, P., Bayly, C. I. & Kollman, P. A. Application of RESP charges to calculate conformational energies, hydrogen bond energies, and free energies of solvation. J. Am. Chem. Soc. 115, 9620–9631 (1993).

34. Eastman, P. et al. OpenMM 7: Rapid development of high performance algorithms for molecular dynamics. PLOS Comput. Biol. 13, e1005659 (2017).

35. Liu, Y., Yu, Q., Wang, D. & Chen, M. AF3Score: A Score-Only Adaptation of AlphaFold3 for Biomolecular Structure Evaluation. J. Chem. Inf. Model. 65, 8207–8214 (2025).

36. Protenix Team et al. Protenix-v1: Toward High-Accuracy Open-Source Biomolecular Structure Prediction. Preprint at 10.64898/2026.02.05.703733 (2026).

