## Supplementary material for "HighFold-Lasso: Topology-Guided Prediction of Lasso Peptide Structures Using AlphaFold3": Sup

**Supplementary data**

**Supplementary Tables**

**Table S1.** Sequences of Lasso Peptides

| PDB ID | Sequence | Type |
| --- | --- | --- |
| 1Q71 | GGAGHVPEYFVGIGTPISFYG | II |
| 1RPB | CLGIGSCNDFAGCGYAVVCFW | I |
| 2LS1 | CVWGGDCTDFLGCGTAWICV | I |
| 2LTI | GLSQGVEPDIGQTYFEESRINQD | II |
| 2LX6 | GAFVGQPEAVNPLGREIQG | II |
| 2M37 | GLSQGVEPDIGQTYFEESR | II |
| 2M8F | GPTPMVGLDSVSGQYWDQHAPLAD | II |
| 2MFV | GGPLAGEEMGGITT | II |
| 2MLJ | SIGDSGLRESMSSQTYWP | II |
| 2MMT | GGAGHVPEYFVRGDFPISFYG | II |
| 2MMW | GGAGHVPEYFVRGDTPISFYG | II |
| 2MW3 | SLGSSPYNDILGYPALIVIYP | II |
| 2N5C | GFGSKPLDSFGLNFF | II |
| 2N68 | GLSQGVEPDIGQTYFEESRINQD | II |
| 2N6U | GLTQIQALDSVSGQFRDQLG | II |
| 2N6V | GPTPMVGLDSVSGQYWDQHAPLAD | II |
| 3NJW | GLPWGCPSDIPGWNTPWAC | III |
| 5D9E | GTLTPGLPEDFLPGHYM(SME)PG | II |
| 5GVO | GLPIGWWIERPSGWYFPI | II |
| 5JPL | GRPNWGFENDWSCVRVC | IV |
| 5OQZ | GAPSLINSEDNPAFPQRV | II |
| 5TJ1 | GVGFGRPDSILTQEQAKPM | II |
| 5UI6 | GGKGPIFETWVTEGNYYG | II |
| 5UI7 | GSDGPIIEFFNPNGVMHYG | II |
| 5XM4 | GPPGDRIEFGVLAQLPG | II |
| 5ZCN | DGMGEEFIEGLVRDSLYPPAG | II |
| 6AK0 | CLGVGSCVDFAGCGYAVVCFW(DTR) | I |
| 6B5W | GVGFGRPDSILTQEQAKPM | II |
| 6M19 | LVVIVQADWNAPGFF | II |
| 6MW6 | GGVGKIIEYFIGGGVGRYG | II |
| 6POR | GGDGSIAEYFNRPMHIHDWQIMDSGYYG | II |
| 6Q1X | GVLGNDAEGITLLPLCFKPICIPTLPPLTGGHA | IV |
| 6XTH | GSRGWGFEPGVRCLIWCD | IV |
| 6XTI | GGGGRGYEYNKQCLIFC | IV |
| 7BW5 | GPKGDFPDVGDGRILAG | II |
| 7BZ7 | LVVIVQADWNAPGWY | II |
| 7BZ8 | LVAIVQADWNAPGWF | II |
| 7BZ9 | LVVAVQADWNAPGWF | II |
| 7BZA | LVVIVQADWNAPGWF | II |
| 7CU6 | LCVIVQADWNCPGWF | Noncanonical‡ |
| 7EES | GTIDPQNSEEHPVLSRRLEN | II |
| 7JS6 | LLGRSGNDRLILSKN | II |
| 7LCW | GSKYSDTADESSYRW | II |
| 7ZWJ | SKKSKPGDGIRGKGVRG | II |
| 8DYN | GHSVDRIPEYFGPPGLPGPVLFYS | II |
| 8SVB | GGGGPTPEYFLMPIDPAWLQANLPNTGKYN | II |
| 8UKC | GIGSQNPW(5CW)DTAW(5CW)LWD | II |
| 8UKG | WY(PQ4)GW(WWB)ASNDGEGTAII | II |
| 9KAR | GGWGTVPDWFFNMNW | II |
| 9LMD | LTNRGYWLDGPWGAWWF | II |
| 9T02 | GISGGTVDAPAGQGLA | II |
| 9Z8J | AKKSKPGDGKHGRGVKK | V |

‡ 7CU6 contains a noncanonical Cys2–Cys11 ring–loop disulfide linkage and was analyzed separately from the groups in Fig. 4A,B.

A parenthetical CCD identifier and its preceding one-letter symbol denote one modified residue, rather than two residues.

**Table S2.** Sequences of Genome-Mined Lasso Peptides

| ID | Sequence |
| --- | --- |
| 1 | GQGKGSAEDKRRAYN |
| 2 | GKNGFDVQDNTEYRQKTPGVSAL |
| 3 | AANPGGSDDMGFARTGGGDGPPDMTFS |
| 4 | LLQRSGNDRLIFSKN |
| 5 | SADGKGDREVGQGDAANQTKSIPS |
| 6 | AGVNGTKFDGNFAAGQPVPFTGGKPAIFS |
| 7 | GAFNSGEGSSGMGPTFVS |
| 8 | GGSMGIAPDEPNDGGWHNLPRH |
| 9 | QRLANALDVPVAYFYAEEDDLAQMILEYKRRGKSVP |
| 10 | GSASGSSDASGQSFN |
| 11 | ASMNEIAPELVGDKTQRFGG |
| 12 | FGGYDSADGAPWQDSNGNWQEPKGKS |
| 13 | GNAGFNEADDTQYWES |
| 14 | ASGGTLKAESVEFIRYDPL |
| 15 | ASVKGSPVDTVFGEDGSIIGNLTGSLDACTSGDLINCGK |
| 16 | AGPGISIADSFQDDPDEIVHYS |
| 17 | TCEQQVLEFINKLKDEELIVITDLNEGCRLL |
| 18 | GSSFTGCAEGAYNKYYWC |
| 19 | TGFPDQQRDFLDTNNTEHLMGPS |
| 20 | GGSTQASELSNNGECKKNVGNGHNNTPRC |
| 21 | GEQIGAGESLGRFGQMGLSDD |
| 22 | ITNGDTRKDFLFLNGTVTSDGNDLGSKDVCLGTTPKGPNCDPKY |
| 23 | AFGASGANDTIQFNGNTFPGSIIGASGSQNGTINPQ |
| 24 | AGAVGVTDAGIFS |
| 25 | RIAAGSAESGKGGGTDNAKTPRS |
| 26 | NSGAPNSDGQGNGTACFGDFCSDPRDP |
| 27 | GMFYASCKEYWSRRLCI |
| 28 | GDAIFQTDGSQGRLEYVTGMADD |
| 29 | ATGPGPKTELNPVFTYEDTGVRAEVVRMGAAETRTRADSEEGQLELIPVQRWEMGGAVR |
| 30 | KSDRVPDGKSGLRGNHS |
| 31 | SIRDGTVSEANNAALKVFPL |
| 32 | GEGLQGSDDTFVFNWGPFHIVIPYGNGS |
| 33 | AGPGIKTPDASQPDVDEVVHYS |
| 34 | ANLVGSLPDALTLVTVSIPGVGNISVPGQGTLPPGVVIVS |
| 35 | AGPGLKYVDSDQNDPDEPDADHYS |
| 36 | TRGVDLEGPVDLQTLMKVRQGGISDED |
| 37 | GSPQGVATELNEAVTYKP |
| 38 | FGCCGRWLDNPFGYGWFDF |
| 39 | AGGPGASDFGIFS |
| 40 | VPQGTAADPTLPGMGILGS |
| 41 | GGPNGVFPDKNGKGGNNGHAHENES |
| 42 | GRPSWGFEADWTCVMFC |
| 43 | GRPGFQSELGVIAQPLGIEAE |
| 44 | NGLAPSEQFDGDWVQIGCKWYVPGHGEELS |
| 45 | LVSAPNNEWIGAYFQILA |
| 46 | AGPGTSIPDAVQPDPDETMHYS |
| 47 | GQAIDWDPVEKGRLWSIDRAKTEMVASFISDGD |
| 48 | GREQRSEKNEAKFRESSS |
| 49 | GEALGARYEGGFPPYEWYLGPS |
| 50 | GLGVEGPIDLATLQRTYNAGIQTDD |
| 51 | ATSGSNTDGMGGNMAPPS |
| 52 | GSILGLLVEDVTAARYDPT |
| 53 | GWGSIGSGDFIFQVVKGEWGTQDGCTGGLDQVFCTGS |
| 54 | AGPGSAIPDGVQPDPDEMVHYS |
| 55 | RNVSGTKTDKAFPAGTPINQLTFST |
| 56 | LGGGAAGTDFGSEISAP |
| 57 | GGQGVLIDFISGEVVAGLSDD |
| 58 | SGRRNNPSESMSSSNASSHVPGTLNS |
| 59 | GQQNFNDADTGGTNYFESAS |
| 60 | VSGRNAIRDTLFFNGQPTNVQSDDSLDLLTTSNGSPTTRPAGQ |
| 61 | RGTQYQDSADDTQYKT |
| 62 | GDDGLPIESGGRMLRPGLTED |
| 63 | KFADIAERSGRAQDKEDGGGLEELRKEGYM |
| 64 | GDIDLPDVEFGDTRYEQP |
| 65 | GGAPSGDFDDPYMPKPWTYNW |
| 66 | AGPGTSTPDAFQPDPDEDVHFDPMDS |
| 67 | NDGNVIGWEEEYGQKPSV |
| 68 | IEFALKRGELRKDLLDYLSSVLDRQLIK |
| 69 | TGGRTTRDGGRGRRRH |
| 70 | NGVLENADVKGGPNGTAECPAGSTDPRCLS |
| 71 | GPGGAIESFGGAPLGGISDA |
| 72 | IEMALQNEELKLDLMKYIEGLVKKEEVHS |
| 73 | LRPGRGADGETIWPDCTS |
| 74 | RGLDGSGPDGCGKAVVFTDACS |
| 75 | GSRLAFEDAWFGVAGTDGLIGPKCEPGQTTWACGS |
| 76 | VKNTGPKDVLGGKQLL |
| 77 | AGPGIATPDAVQPDPTEVIHFS |
| 78 | LFGSGSRNDFFIFTAPPNGINVQPNQPVPATYGSQDGVLSPSPTKVN |
| 79 | AGDIGEPEPDVGQLFE |
| 80 | GGTASGSENKRKVYSSTIG |
| 81 | NGPGNANADCFDVSDGKSETHEGRSNASCLQGS |
| 82 | WSSPAQPDLPGGRAGPGGRAAHSRLNGRGMRNAAASAASLWGVAGAEPPQKGSVETRGLGGRGRHSPPYAKRGDVGR |
| 83 | GSAGNNNADDTQYWE |
| 84 | ANGSINLADVFILSNLPGEQPDPITADGQGSIDIEIDVEGAI |
| 85 | AFGPCQEQDWFDGTAWIC |
| 86 | NDKIGDSEDAVTEAAADTGASITGSIF |
| 87 | GRQAFNDSDKSNYFEG |
| 88 | SGTSGTSENKPCGSGQGPPSKKCV |
| 89 | GGTMGPYCDVFVGQLYPVNDTVMNGNCWKDMP |
| 90 | SGIVGSPEDFLPGRHMPA |
| 91 | FSGNQTRKDFVFFNGDNNAGLSGPVNLNGSLGSIDHFDTPCPENPSRCK |
| 92 | SGSPYVGTEDGLYAPLES |
| 93 | TNSMTGMADGGGGAMNRTG |
| 94 | GAAVGSQMDGIAAQKT |
| 95 | GLWFGKFTDIFGGRAVFQIVINY |
| 96 | TGSLNGQEDLVENGVCTMIDINNINNMYNMC |
| 97 | GNGGGGFDFGSEVSA |
| 98 | GVTPGIADLEFGQQLSMGLSDD |
| 99 | GTGELGPLDIQTFQRTYNAGIQTDD |
| 100 | GPSSGTSDGKSAGAMAMG |

**Table S3.** RMSD and pLDDT of AlphaFold3 predictions for peptides in the lasso peptide monomer dataset.

| PDB ID | RMSD_Cα_ | pLDDT |
| --- | --- | --- |
| 1Q71 | 1.065 | 91.875 |
| 1RPB | 2.458 | 87.938 |
| 2LS1 | 0.600 | 95.351 |
| 2LTI | 6.864 | 87.571 |
| 2LX6 | 3.929 | 96.369 |
| 2M37 | 2.171 | 87.451 |
| 2M8F | 5.847 | 86.675 |
| 2MFV | 0.411 | 95.274 |
| 2MLJ | 0.903 | 89.439 |
| 2MMT | 3.063 | 88.292 |
| 2MMW | 1.524 | 90.832 |
| 2MW3 | 0.908 | 90.862 |
| 2N5C | 0.790 | 87.088 |
| 2N68 | 4.158 | 87.571 |
| 2N6U | 0.964 | 91.814 |
| 2N6V | 1.890 | 86.675 |
| 3NJW | 0.556 | 87.425 |
| 5D9E | 0.710 | 96.208 |
| 5GVO | 0.710 | 84.889 |
| 5JPL | 0.661 | 89.097 |
| 5OQZ | 0.556 | 93.571 |
| 5TJ1 | 2.146 | 82.242 |
| 5UI6 | 1.047 | 91.389 |
| 5UI7 | 1.082 | 85.643 |
| 5XM4 | 0.921 | 93.746 |
| 5ZCN | 8.127 | 88.134 |
| 6AK0 | 2.047 | 90.270 |
| 6B5W | 2.251 | 82.242 |
| 6M19 | 1.205 | 77.555 |
| 6MW6 | 1.190 | 85.516 |
| 6POR | 1.871 | 86.459 |
| 6Q1X | 5.169 | 73.342 |
| 6XTH | 0.828 | 83.761 |
| 6XTI | 1.187 | 91.646 |
| 7BW5 | 1.294 | 92.563 |
| 7BZ7 | 2.374 | 74.210 |
| 7BZ8 | 1.948 | 77.106 |
| 7BZ9 | 1.499 | 75.950 |
| 7BZA | 2.302 | 79.832 |
| 7CU6 | 3.350 | 82.063 |
| 7EES | 5.352 | 80.172 |
| 7JS6 | 2.153 | 79.443 |
| 7LCW | 1.492 | 89.534 |
| 7ZWJ | 7.333 | 83.069 |
| 8DYN | 3.716 | 87.000 |
| 8SVB | 8.872 | 75.912 |
| 8UKC | 4.752 | 73.664 |
| 8UKG | 4.224 | 76.190 |
| 9KAR | 4.884 | 85.327 |
| 9LMD | 4.983 | 69.869 |
| 9T02 | 4.907 | 88.093 |
| 9Z8J | 5.971 | 75.113 |

**Table S4.** Ring, ring+loop, and full-structure RMSDs of LassoPred predictions.

| PDB ID | Ring_Cα_ | Ring_all-atom_ | Ring+Loop_Cα_ | Ring+Loop_all-atom_ | Full_Cα_ | Full_all-atom_ |
| --- | --- | --- | --- | --- | --- | --- |
| 1Q71 | 1.226 | 2.095 | 2.752 | 3.961 | 2.719 | 3.878 |
| 1RPB | 0.815 | 1.898 | 1.788 | 2.807 | 5.278 | 6.798 |
| 2LS1 | 1.112 | 3.157 | 2.099 | 4.141 | 3.277 | 5.591 |
| 2LTI | 1.052 | 2.923 | 3.531 | 5.795 | 6.052 | 8.029 |
| 2LX6 | 1.368 | 3.008 | 2.570 | 4.557 | 2.881 | 4.653 |
| 2M37 | 1.640 | 3.173 | 2.415 | 3.574 | 2.700 | 4.232 |
| 2M8F | 1.840 | 4.574 | 3.773 | 5.909 | 6.550 | 8.545 |
| 2MFV | 0.221 | 0.872 | 0.201 | 1.104 | 2.370 | 3.169 |
| 2MLJ | 2.036 | 4.847 | 2.476 | 5.229 | 2.596 | 5.424 |
| 2MMT | 0.836 | 1.840 | 2.475 | 4.935 | 2.587 | 4.945 |
| 2MMW | 1.148 | 1.914 | 2.649 | 5.090 | 2.799 | 5.145 |
| 2MW3 | 1.966 | 3.146 | 2.148 | 3.054 | 3.502 | 4.345 |
| 2N5C | 1.724 | 4.886 | 2.910 | 6.504 | 3.276 | 6.452 |
| 2N68 | 1.585 | 2.983 | 2.227 | 3.546 | 5.183 | 6.598 |
| 2N6U | 1.648 | 4.399 | 2.017 | 4.353 | 3.667 | 5.655 |
| 2N6V | 1.634 | 4.523 | 3.605 | 5.802 | 6.184 | 7.524 |
| 3NJW | 1.569 | 2.651 | 3.303 | 4.579 | 4.233 | 5.436 |
| 5D9E | 0.100 | 1.365 | 0.086 | 1.203 | 1.003 | 2.087 |
| 5GVO | 1.741 | 3.770 | 1.819 | 4.037 | 1.996 | 4.539 |
| 5JPL | 0.117 | 2.571 | 0.106 | 2.643 | 2.842 | 4.344 |
| 5OQZ | 0.090 | 0.925 | 0.083 | 0.835 | 0.277 | 1.432 |
| 5TJ1 | 1.019 | 3.553 | 1.962 | 4.053 | 3.048 | 4.927 |
| 5UI6 | 1.014 | 2.531 | 2.405 | 4.101 | 2.296 | 4.147 |
| 5UI7 | 1.342 | 2.005 | 3.219 | 4.936 | 3.191 | 4.977 |
| 5XM4 | 1.489 | 3.170 | 2.312 | 4.342 | 3.074 | 4.840 |
| 5ZCN | 1.770 | 4.233 | 2.010 | 4.018 | 3.524 | 4.976 |
| 6AK0 | 0.917 | 2.395 | 1.602 | 2.718 | 4.151 | 5.425 |
| 6B5W | 1.106 | 4.202 | 2.551 | 4.763 | 3.097 | 5.192 |
| 6M19 | 2.000 | 3.420 | 3.137 | 4.994 | 3.412 | 5.074 |
| 6MW6 | 1.059 | 2.346 | 3.375 | 5.390 | 3.231 | 5.235 |
| 6POR | 1.038 | 1.431 | 4.225 | 6.430 | 4.115 | 6.351 |
| 6Q1X | 0.268 | 1.504 | 0.239 | 1.792 | 11.762 | 11.760 |
| 6XTH | 1.478 | 3.852 | 2.221 | 3.924 | 4.601 | 6.098 |
| 6XTI | 1.643 | 2.347 | 1.743 | 2.883 | 3.787 | 4.846 |
| 7BW5 | 1.500 | 3.253 | 1.450 | 3.211 | 2.625 | 4.730 |
| 7BZ7 | 2.027 | 3.790 | 3.229 | 5.216 | 3.493 | 5.223 |
| 7BZ8 | 2.165 | 4.001 | 3.134 | 4.730 | 3.455 | 4.984 |
| 7BZ9 | 1.683 | 3.065 | 2.960 | 4.993 | 3.085 | 5.020 |
| 7BZA | 2.071 | 3.361 | 3.124 | 4.974 | 3.251 | 5.035 |
| 7CU6 | 1.634 | 3.566 | 3.225 | 5.157 | 3.506 | 5.300 |
| 7EES | 1.843 | 3.569 | 2.131 | 3.704 | 2.906 | 4.443 |
| 7JS6 | 1.704 | 4.230 | 2.528 | 4.814 | 2.439 | 4.788 |
| 7LCW | 0.076 | 1.664 | 0.071 | 1.563 | 0.213 | 2.289 |
| 7ZWJ | 1.042 | 2.640 | 1.791 | 3.940 | 2.312 | 4.529 |
| 8DYN | 2.016 | 4.411 | 3.057 | 4.738 | 2.983 | 4.749 |
| 8SVB | 0.763 | 1.474 | 7.628 | 8.971 | 7.592 | 9.116 |
| 8UKC | 2.176 | 4.320 | 2.885 | 4.828 | 2.761 | 4.895 |
| 8UKG | 1.934 | 4.204 | 2.747 | 5.019 | 3.131 | 5.260 |
| 9KAR | 1.166 | 3.605 | 1.260 | 3.939 | 2.146 | 4.141 |
| 9LMD | 2.070 | 5.380 | 4.421 | 6.664 | 4.977 | 7.492 |
| 9T02 | 1.469 | 3.041 | 3.061 | 4.113 | 3.491 | 4.617 |
| 9Z8J | 1.048 | 4.075 | 2.310 | 4.971 | 2.634 | 5.609 |

**Table S5.** Ring, ring+loop, and full-structure RMSDs of HighFold-Lasso predictions.

| PDB ID | Ring_Cα_ | Ring_all-atom_ | Ring+Loop_Cα_ | Ring+Loop_all-atom_ | Full_Cα_ | Full_all-atom_ |
| --- | --- | --- | --- | --- | --- | --- |
| 1Q71 | 0.492 | 0.826 | 3.576 | 4.796 | 3.519 | 4.671 |
| 1RPB | 1.149 | 1.709 | 1.253 | 1.823 | 1.430 | 2.751 |
| 2LS1 | 0.718 | 1.180 | 0.706 | 1.203 | 0.717 | 1.235 |
| 2LTI | 1.641 | 2.522 | 2.285 | 3.501 | 2.540 | 3.907 |
| 2LX6 | 1.045 | 2.159 | 1.551 | 2.697 | 1.814 | 2.733 |
| 2M37 | 1.234 | 2.244 | 1.360 | 2.154 | 1.607 | 3.245 |
| 2M8F | 1.353 | 2.287 | 2.205 | 3.163 | 4.402 | 5.791 |
| 2MFV | 0.454 | 0.764 | 0.443 | 0.910 | 0.569 | 1.040 |
| 2MLJ | 0.602 | 1.331 | 0.711 | 1.234 | 0.699 | 1.188 |
| 2MMT | 0.838 | 1.157 | 3.214 | 5.521 | 3.094 | 5.446 |
| 2MMW | 0.618 | 0.963 | 3.096 | 4.174 | 2.997 | 4.062 |
| 2MW3 | 0.901 | 1.646 | 0.837 | 1.422 | 1.449 | 1.970 |
| 2N5C | 0.694 | 1.857 | 0.893 | 1.740 | 1.165 | 1.879 |
| 2N68 | 1.131 | 2.323 | 1.077 | 2.280 | 3.926 | 5.473 |
| 2N6U | 0.601 | 1.317 | 1.437 | 2.197 | 1.333 | 2.053 |
| 2N6V | 0.988 | 1.981 | 1.891 | 2.804 | 4.432 | 5.410 |
| 3NJW | 0.883 | 1.348 | 0.968 | 1.644 | 1.047 | 1.987 |
| 5D9E | 0.195 | 0.286 | 0.307 | 0.412 | 0.429 | 0.861 |
| 5GVO | 0.614 | 1.291 | 0.717 | 1.375 | 0.798 | 1.572 |
| 5JPL | 0.883 | 1.713 | 0.776 | 1.572 | 1.449 | 2.015 |
| 5OQZ | 0.323 | 0.753 | 1.016 | 1.405 | 0.964 | 1.329 |
| 5TJ1 | 0.539 | 1.231 | 2.617 | 3.839 | 2.824 | 4.334 |
| 5UI6 | 0.955 | 1.817 | 1.938 | 3.088 | 1.864 | 2.999 |
| 5UI7 | 0.623 | 1.112 | 2.081 | 2.792 | 2.070 | 2.747 |
| 5XM4 | 0.603 | 1.257 | 0.727 | 1.252 | 0.949 | 1.420 |
| 5ZCN | 1.572 | 2.497 | 1.685 | 3.645 | 2.306 | 4.243 |
| 6AK0 | 0.954 | 1.837 | 1.675 | 2.433 | 3.478 | 4.969 |
| 6B5W | 0.764 | 2.350 | 2.702 | 4.003 | 2.741 | 4.101 |
| 6M19 | 1.711 | 2.798 | 2.020 | 3.120 | 2.451 | 4.126 |
| 6MW6 | 0.504 | 1.088 | 0.838 | 1.815 | 0.837 | 1.742 |
| 6POR | 1.165 | 1.556 | 2.566 | 4.415 | 2.539 | 4.376 |
| 6Q1X | 0.972 | 1.791 | 4.535 | 5.534 | 6.387 | 7.143 |
| 6XTH | 0.655 | 1.506 | 0.701 | 1.512 | 0.903 | 1.751 |
| 6XTI | 1.063 | 1.689 | 1.222 | 2.197 | 1.515 | 2.591 |
| 7BW5 | 0.773 | 1.180 | 0.951 | 1.486 | 1.443 | 2.300 |
| 7BZ7 | 0.872 | 1.937 | 1.919 | 2.852 | 2.350 | 3.973 |
| 7BZ8 | 1.659 | 2.310 | 2.072 | 2.950 | 2.017 | 3.001 |
| 7BZ9 | 1.016 | 2.104 | 1.698 | 2.581 | 2.227 | 4.116 |
| 7BZA | 1.881 | 3.058 | 2.316 | 3.766 | 2.359 | 3.888 |
| 7CU6 | 0.832 | 1.976 | 0.977 | 2.409 | 1.287 | 2.952 |
| 7EES | 2.052 | 3.137 | 2.362 | 3.701 | 3.856 | 4.792 |
| 7JS6 | 1.414 | 4.175 | 1.979 | 3.907 | 1.881 | 3.955 |
| 7LCW | 0.649 | 1.362 | 0.615 | 1.276 | 0.757 | 1.653 |
| 7ZWJ | 1.364 | 2.918 | 1.496 | 3.039 | 2.669 | 3.931 |
| 8DYN | 1.972 | 3.309 | 4.864 | 5.896 | 4.831 | 5.782 |
| 8SVB | 1.431 | 1.612 | 8.036 | 8.496 | 8.058 | 8.616 |
| 8UKC | 1.926 | 3.656 | 2.564 | 3.884 | 2.465 | 4.000 |
| 8UKG | 1.228 | 4.550 | 1.525 | 4.729 | 1.986 | 5.144 |
| 9KAR | 1.757 | 2.619 | 2.571 | 3.903 | 3.391 | 4.406 |
| 9LMD | 1.694 | 3.517 | 3.202 | 4.546 | 3.599 | 5.669 |
| 9T02 | 1.154 | 2.194 | 3.413 | 3.877 | 4.056 | 4.608 |
| 9Z8J | 1.677 | 2.961 | 2.773 | 4.206 | 3.035 | 4.761 |

**Table S6.** RMSD and pLDDT of peptides in the lasso peptide monomer dataset.

| PDB ID | LassoPred pLDDT | LassoPred RMSD _Cα_ | HF-Lasso pLDDT | HF-Lasso RMSD _Cα_ |
| --- | --- | --- | --- | --- |
| 1Q71 | 90.12191 | 2.71867 | 94.78966 | 3.51916 |
| 1RPB | 86.05092 | 5.27822 | 91.15611 | 1.43039 |
| 2LS1 | 93.21798 | 3.27745 | 95.06778 | 0.71677 |
| 2LTI | 80.55763 | 6.05247 | 90.32881 | 2.54048 |
| 2LX6 | 93.45012 | 2.88128 | 96.07315 | 1.81419 |
| 2M37 | 91.02631 | 2.70028 | 94.09936 | 1.60672 |
| 2M8F | 85.62899 | 6.55033 | 87.07334 | 4.40163 |
| 2MFV | 94.13418 | 2.36955 | 95.33647 | 0.56874 |
| 2MLJ | 86.53614 | 2.59554 | 88.09071 | 0.69917 |
| 2MMT | 87.43791 | 2.58706 | 90.68041 | 3.09388 |
| 2MMW | 88.43109 | 2.79940 | 89.01187 | 2.99708 |
| 2MW3 | 91.17044 | 3.50227 | 93.15533 | 1.44924 |
| 2N5C | 89.22481 | 3.27641 | 90.65501 | 1.16512 |
| 2N68 | 89.87061 | 5.18342 | 85.66547 | 3.92573 |
| 2N6U | 89.58524 | 3.66653 | 92.62698 | 1.33276 |
| 2N6V | 85.78253 | 6.18408 | 88.41131 | 4.43210 |
| 3NJW | 85.23983 | 4.23288 | 93.36797 | 1.04726 |
| 5D9E | 96.94222 | 1.00340 | 97.42131 | 0.42876 |
| 5GVO | 88.68632 | 1.99567 | 86.05735 | 0.79753 |
| 5JPL | 92.46799 | 2.84155 | 88.22884 | 1.44912 |
| 5OQZ | 91.01672 | 0.27676 | 91.30529 | 0.96409 |
| 5TJ1 | 88.23164 | 3.04791 | 91.21844 | 2.82405 |
| 5UI6 | 88.04288 | 2.29551 | 91.96533 | 1.86392 |
| 5UI7 | 87.47838 | 3.19142 | 88.56997 | 2.06954 |
| 5XM4 | 93.18089 | 3.07444 | 97.56448 | 0.94871 |
| 5ZCN | 79.46712 | 3.52382 | 82.88859 | 2.30556 |
| 6AK0 | 87.44295 | 4.15134 | 89.47581 | 3.47790 |
| 6B5W | 87.17212 | 3.09742 | 91.44867 | 2.74057 |
| 6M19 | 77.03214 | 3.41199 | 80.87028 | 2.45121 |
| 6MW6 | 84.70519 | 3.23103 | 86.69229 | 0.83724 |
| 6POR | 84.15232 | 4.11526 | 82.71943 | 2.53924 |
| 6Q1X | 68.51390 | 11.76189 | 71.25608 | 6.38749 |
| 6XTH | 90.03407 | 4.60072 | 90.31013 | 0.90267 |
| 6XTI | 90.35106 | 3.78680 | 87.46527 | 1.51486 |
| 7BW5 | 92.54542 | 2.62455 | 96.86388 | 1.44331 |
| 7BZ7 | 70.97087 | 3.49295 | 78.72679 | 2.34966 |
| 7BZ8 | 73.01306 | 3.45461 | 81.15074 | 2.01700 |
| 7BZ9 | 73.24893 | 3.08488 | 81.84047 | 2.22659 |
| 7BZA | 75.14239 | 3.25100 | 79.28641 | 2.35919 |
| 7CU6 | 80.05897 | 3.50623 | 82.32360 | 1.28705 |
| 7EES | 89.06718 | 2.90625 | 88.38795 | 3.85636 |
| 7JS6 | 77.71410 | 2.43906 | 76.40343 | 1.88064 |
| 7LCW | 87.39228 | 0.21253 | 85.72999 | 0.75684 |
| 7ZWJ | 89.35514 | 2.31249 | 86.81431 | 2.66923 |
| 8DYN | 82.38394 | 2.98313 | 84.84764 | 4.83097 |
| 8SVB | 77.19473 | 7.59221 | 75.43473 | 8.05759 |
| 8UKC | 84.68673 | 2.76113 | 86.85734 | 2.46534 |
| 8UKG | 74.90420 | 3.13142 | 86.81832 | 1.98572 |
| 9KAR | 85.53034 | 2.14606 | 80.80519 | 3.39146 |
| 9LMD | 69.85797 | 4.97653 | 78.68301 | 3.59867 |
| 9T02 | 94.38515 | 3.49079 | 92.82926 | 4.05601 |
| 9Z8J | 77.60207 | 2.63357 | 81.09744 | 3.03547 |

**Table S7.** RMSD_Cα_ between HighFold-Lasso input backbones and output structures.

| **PDB ID** | **Ring** | **Ring+Loop** | **Full** |
| --- | --- | --- | --- |
| 1Q71 | 2.728 | 3.187 | 3.151 |
| 1RPB | 2.942 | 3.621 | 4.869 |
| 2LS1 | 2.592 | 2.791 | 3.427 |
| 2LTI | 2.884 | 3.831 | 5.692 |
| 2LX6 | 1.645 | 2.404 | 2.377 |
| 2M37 | 1.518 | 1.775 | 1.976 |
| 2M8F | 4.480 | 4.613 | 5.175 |
| 2MFV | 1.801 | 1.665 | 2.128 |
| 2MLJ | 2.412 | 2.529 | 2.568 |
| 2MMT | 2.580 | 3.278 | 3.240 |
| 2MMW | 1.881 | 2.968 | 2.945 |
| 2MW3 | 2.858 | 2.553 | 2.812 |
| 2N5C | 3.451 | 3.351 | 3.453 |
| 2N68 | 2.162 | 2.427 | 2.623 |
| 2N6U | 2.801 | 2.940 | 3.734 |
| 2N6V | 4.541 | 4.647 | 5.203 |
| 3NJW | 3.896 | 4.075 | 4.467 |
| 5D9E | 0.631 | 0.764 | 1.079 |
| 5GVO | 1.992 | 1.814 | 1.908 |
| 5JPL | 2.332 | 2.078 | 3.191 |
| 5OQZ | 0.936 | 1.010 | 0.980 |
| 5TJ1 | 2.016 | 2.478 | 2.519 |
| 5UI6 | 1.643 | 1.703 | 1.643 |
| 5UI7 | 2.024 | 2.666 | 2.600 |
| 5XM4 | 4.010 | 3.553 | 3.332 |
| 5ZCN | 2.935 | 2.821 | 2.944 |
| 6AK0 | 3.662 | 3.555 | 4.478 |
| 6B5W | 2.037 | 2.438 | 2.529 |
| 6M19 | 2.651 | 2.699 | 2.802 |
| 6MW6 | 2.303 | 3.421 | 3.300 |
| 6POR | 2.244 | 4.432 | 4.338 |
| 6Q1X | 12.110 | 11.335 | 10.538 |
| 6XTH | 3.662 | 3.464 | 4.421 |
| 6XTI | 3.054 | 2.612 | 3.516 |
| 7BW5 | 1.584 | 1.552 | 1.872 |
| 7BZ7 | 4.223 | 4.331 | 4.674 |
| 7BZ8 | 3.497 | 3.597 | 3.567 |
| 7BZ9 | 3.012 | 3.317 | 3.736 |
| 7BZA | 2.850 | 2.623 | 2.774 |
| 7CU6 | 2.917 | 3.192 | 3.134 |
| 7EES | 2.597 | 2.722 | 2.725 |
| 7JS6 | 2.051 | 2.092 | 1.955 |
| 7LCW | 0.734 | 0.655 | 0.671 |
| 7ZWJ | 2.533 | 2.969 | 3.951 |
| 8DYN | 2.705 | 4.055 | 4.034 |
| 8SVB | 2.476 | 3.963 | 3.836 |
| 8UKC | 1.574 | 1.688 | 1.646 |
| 8UKG | 2.238 | 2.301 | 2.395 |
| 9KAR | 2.788 | 2.701 | 2.868 |
| 9LMD | 3.641 | 3.377 | 3.333 |
| 9T02 | 2.314 | 2.243 | 2.392 |
| 9Z8J | 2.253 | 2.384 | 2.609 |
